# Generation and validation of a *floxed FosB* mouse line

**DOI:** 10.1101/179309

**Authors:** Yoshinori N. Ohnishi, Andrew L. Eagle, Yoko H. Ohnishi, Michael E. Cahill, Alexis J. Wirtz, Alfred J. Robison, Eric J. Nestler

**Affiliations:** Department of Pharmacology, Kurume University School of Medicine, Kurume, Fukuoka, Japan; Department of Neuroscience and Friedman Brain Institute, Icahn School of Medicine at Mount Sinai, New York, NY, USA; Department of Physiology, Michigan State University, East Lansing, MI, USA; Department of Comparative Biosciences, University of Wisconsin at Madison, Madison, WI, USA

## Abstract

Expression of the *FosB* gene has been studied extensively in many fields using a variety of tools. However, previous techniques have had a variety of caveats, from potential off-target effects (e.g., overexpression of FosB, ΔFosB, or a dominant negative mutant of JunD, termed ΔJunD) or confounding developmental effects (e.g., the constitutive *FosB* knockout mouse). Therefore, we sought to create a *floxed FosB* mouse line that will allow true silencing of the *FosB* gene with both spatial and temporal control. Here, we detail the cloning strategy, production, and validation of the *floxed FosB* mouse. We demonstrate methodology for breeding and genotyping, and show that viral-mediated expression of Cre recombinase in a targeted, discrete brain region ablates expression of the *FosB* gene in *floxed* but not wild type mice. Thus, the *floxed FosB* mouse presented here represents an important new tool for the continued investigation of this critical gene.

## Introduction

*FosB* is an activity-dependent immediate early gene whose expression is enriched in the brain. It encodes transcription factors that combine with Jun-family proteins to form AP-1 complexes that bind DNA and regulate the transcription of a host of gene targets. Expression of the *FosB* gene has been studied extensively in the fields of addiction (Robison and Nestler, 2011), mood disorders (Manning et al., 2017), natural reward (Teegarden et al., 2008; Pitchers et al., 2010), learning (Eagle et al., 2015), Parkinson’s disease and related syndromes (Dietz et al., 2014; Feyder et al., 2016), epilepsy (Giordano et al., 2015), bone density (Rowe et al., 2012), and cataracts (Kelz et al., 2000). Its role in long-term behavioral adaptations underlying both maladaptive and normal forms of learning, reward, and other functions has been speculated to arise from the unique ability of its protein products to modulate long-lasting changes in gene expression.

In the rodent brain, *FosB* gene products are spliced to produce two major mRNA variants: a full-length FosB variant that results in a 338-amino acid protein, and an alternative variant that introduces a premature stop codon to produce a 237-amino acid c-terminal truncation termed ΔFosB (Nestler, 2015). The C-terminal region of FosB missing in the ΔFosB isoform contains a portion of the putative transactivation domain as well as two degron domains that target the full-length protein for proteosomal degradation (Carle et al., 2007). Thus, the ΔFosB isoform is uniquely stable, with a half-life *in vivo* of approximately eight days (Ulery-Reynolds et al., 2009), making it particularly well-suited to mediate long-term changes in gene expression.

A variety of tools have been used to interrogate the function of the *FosB* gene in the brain. Multiple mouse lines have been used to overexpress the ΔFosB isoform in select neuronal populations, establishing that ΔFosB expression in D1-type medium spiny neurons of the nucleus accumbens (NAc) is sufficient to increase the rewarding properties of multiple drugs (Kelz et al., 1999; Zachariou et al., 2006) as well as to mediate stress resilience (Vialou et al., 2010; Ohnishi et al., 2015). Viral vectors overexpressing ΔFosB in all NAc neurons have been used to confirm these results. In order to establish the necessity of ΔFosB expression for these processes, most studies have relied on overexpression of mutant forms of the requisite binding partners, Jun family proteins termed ΔJunD or Δc-Jun, which lack their N-terminal transactivation domains and presumably act as dominant negative antagonists *in vivo* (Peakman et al., 2003; Zachariou et al., 2006; Vialou et al., 2010). The use of transgenic mice expressing these inhibitors, and of Cre recombinase-dependent viral vectors, has allowed cell-type specific inhibition of ΔFosB-dependent transcriptional activity, but questions remain as to whether ΔJunD has off-target effects and the extent to which it is able to inhibit the entire cellular pool of ΔFosB.

*FosB* knockout mouse lines have been produced as well (Brown et al., 1996; Yutsudo et al., 2013). Consistent with the overexpression studies cited above, one of the lines shows increased susceptibility to depression-like behaviors (Yutsudo et al., 2013). On the other hand, a different knockout line shows enhanced, not reduced, sensitivity to cocaine (Hiroi et al., 1997), the opposite of what has been observed with the more selective transgenic and viral tools. The whole-body nature of the knockout animal by definition suggests that any behavioral effects in the adult may arise from altered development. Indeed, the *FosB* knockout mouse lines show either defects in maternal behavior (Brown et al., 1996) or a malformed dentate gyrus (Yutsudo et al., 2013). As well, the knockout mice cannot provide information on the site of action of *FosB* gene products within any given brain region or cell type. Therefore, we sought to create a *floxed FosB* mouse line. Such a line, when mated with Cre-recombinase expressing mouse lines or in combination with viral vectors encoding Cre, will allow true knockout of the *FosB* gene with both spatial and temporal control, and without the caveats of non-specific inhibition of other Fos-family proteins or the constitutive knockout.

### Construction of targeting vector

We cloned the *FRT*-neo cassette-*FRT-loxP* region from a neo expressing DNA vector and fragments of the *FosB* gene between the NheI site of intron 1 and the ScaI site in intron 3 (including a *loxP* sequence at the ScaI site) into the pENTR/D-TOPO vector (Invitrogen, Massachusetts, USA). After DNA sequencing of the cloning products, we combined both products into the *FosB* gene cloning vector with tyrosine kinase expressing cassettes at both ends of the *FosB* gene construct (Figure 1A). The *FRT*-neo cassette-*FRT-loxP* region was then inserted into the NheI site of intron 1 of the *FosB* gene. The second *lox*P was inserted at the ScaI site in intron 3. These steps made use of the following cloning primers:

**Figure 1:**
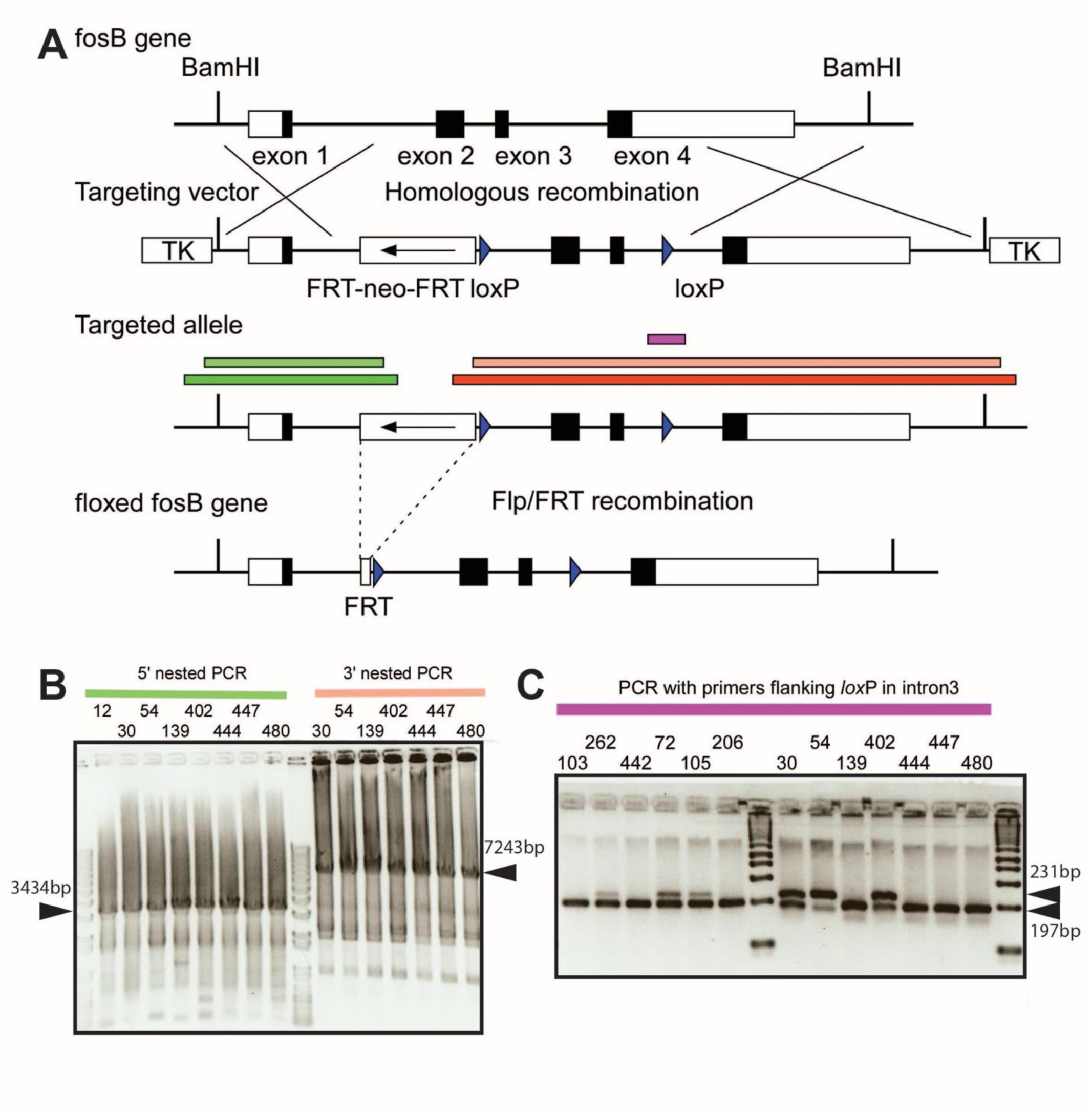
*FosB* targeting and construct cloning strategy. A) Global map of the *FosB* gene and targeting gene construct. Colored bars represent products of nested PCR or genotyping PCR. B) 1% agarose gel of DNA products of nested genomic PCR of various ES cells (numbered at top) between the 5’ outside region of the targeting construct area and its inside region (left) and nested genomic PCR between the 3’ outside region of the targeting construct area and its inside region (right). Black arrows indicate expected bands with given correct homologous recombination. C) 4% agarose gel of DNA products of PCR around the second *loxP* region. Successful homologous recombination around second *loxP* region results in two bands, 231 bp and 197 bp, indicated by black arrows.

<u>Cloning primers for neo cassette</u>

FRTneoUT; 5’-caccGCTAGCctgccataactagtccttgGAAGTTCCTATTCTCTAGAAAGTATAGGA-ACttcGATGAGCTTTATCCAAACC-3’,

*loxP*FRTneoL; 5’-CATATGataacttcgtatagcatacattatacgaagttatGAAGTTCCTATACTTTCTAGA-GAATAGGAACttcGGCGCGCCGCTCGCGAAAGCTTGGGC-3’

<u>Cloning primers for second *loxP* site</u>

FLOXUT; 5’-caccCATATGgAtagcCTACGGAGAGGCAGCCAGGTGGTCTCTAAAAGGTC-3’,

FLOXL; 5’-AGTACTataacttcgtatagcatacattatacgaagttatCTGCCTTAAAGGGCAGAAGGGGGC-ATTGTAGCTCACTGAGTGAAGGAGATTG-3’

### Genotyping of ES cells

We performed nested genomic PCR for confirmation of homologous recombination around the *FosB* gene in ES cells (Figure 1A and B) using Phire Taq (NEB, Massachusetts, USA). Additionally, we examined whether the homologous recombination covered the *loxP* site in intron 3 by PCR with a separate primer set (Figure 1A and C). These steps made use of the following primers:

<u>1^st^ PCR for 5’ site (size 5900 bp)</u>

5FB1stU; 5’-AGAGGTAGAGGTCATCCTG-3’

3FB1stL; 5’-GATTGCACGCAGGTTCTCCG-3’

<u>Nested PCR for 5’ site (size 3434 bp)</u>

5FB2ndU; 5’-CCAGGACTTCACACATGCT-3’

5FB2ndL; 5’-AGAGCGAGGGAAGCGTCTACCTA-3’

<u>1st PCR for 3’ site (size 8936 bp)</u>

3FB1stU; 5’-GAGCCACCTTCTTCTCCAA-3’

3FB1stL; 5’-CAAAACCTCGCCTCCAAGT-3’

<u>Nested PCR for 3’ site (size 7243 bp)</u>

3FB2ndU; 5’-TGGCGCGTCCAATCAATTG-3’

3FB2ndL; 5’-TCTCTGCGTTGGAGCAGTA-3’

<u>PCR for *loxP* in intron 3 (size 231 bp or 197 bp)</u>

2ndLOXPU; 5’-CAATGCCCCCTTCTGCCCTTTA-3’

2ndLOXPL; 5’-TGCTACTTGTGCCTCGGTTTCC-3’

### Generation of *floxed FosB* mice

*Floxed FosB* embryonic stem (ES) cell clones with correct homologous recombination were injected into blastocysts prepared from C57BL/6J mice. The *floxed FosB* allele contained two *loxP* sites in intron 1 and 3, and an *FRT*-neo-*FRT* cassette next to *loxP* in intron 1 (Figure 1). After Cre recombination, the *FosB* gene will lose exons 2 and 3 (Figure 2A), which include the N-terminal transactivation domain and basic region and most of the leucine zipper domain, regions with essential roles in DNA binding, transactivation, and dimerization with Jun family proteins. Neomycin-resistant gene cassettes for positive selection were removed by Flp/*FRT*-mediated recombination through breeding with FLPo transgenic mice. This work was conducted with the Mouse Genetics and Gene Targeting (MGGT) CoRE at Mount Sinai.

**Figure 2:**
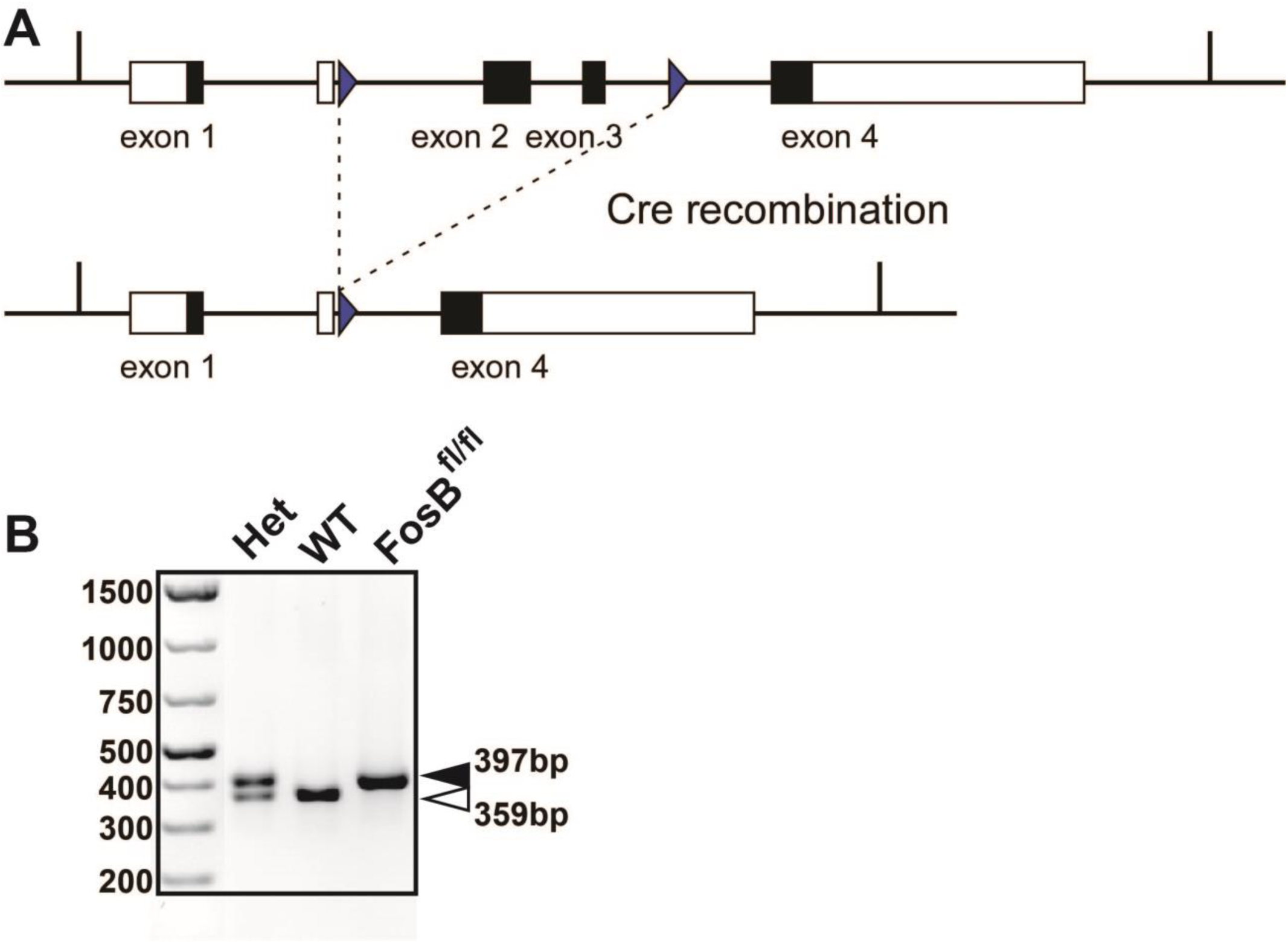
*Floxed FosB* mouse recombination strategy and genotyping. A) Cre-mediated recombination of the *floxed FosB* gene results in removal of exons 2 and 3, functionally knocking out expression of the gene. B) 2% agarose gel of PCR products from tail-derived DNA used to genotype *floxed FosB* mice. Wild type mice have a product of 359 bp (white arrow), *floxed FosB* mice have a product of 397 bp (black arrow), and heterozygous mice have both bands.

### Genotyping and breeding of *floxed FosB* mice

Mice were back-crossed with wild-type C57BL/6J obtained from Jackson Labs (Bar Harbor, ME, USA) for at least four generations. PCR genotyping was performed on tail-derived DNA using the REDExtract-N-Amp kit (Sigma-Aldrich, St. Louis, MO, USA) with the following thermocycler protocol:

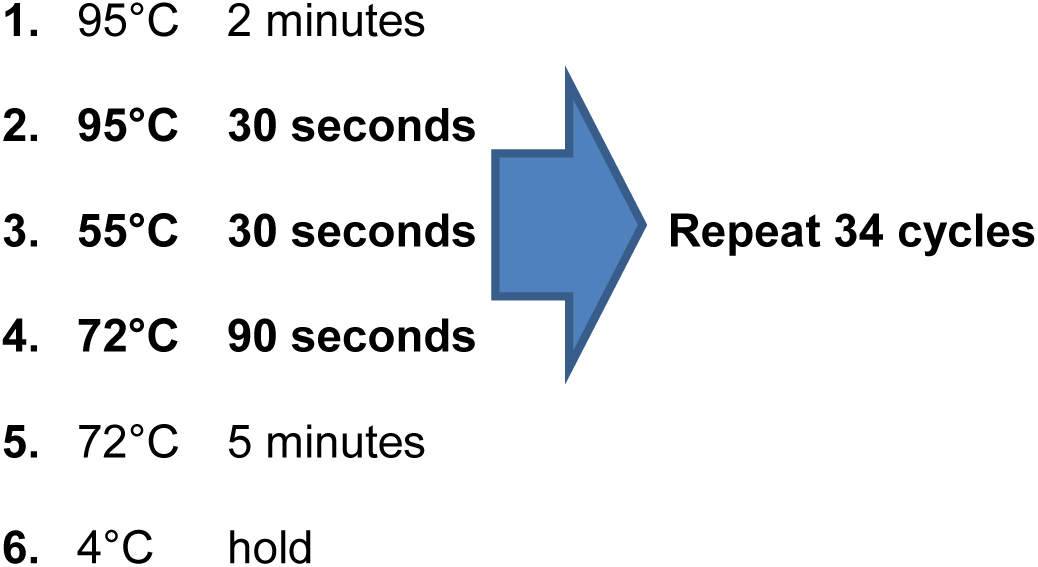

PCR primers were as follows:

FB loxPu sequence; 5’ – GCT GAA GGA GAT GGG TAA CAG – 3’

LIPz sequence; 5’ – AAG CCT GGT GTG ATG GTG A – 3’

LNEo1 sequence; 5’ – AGA GCG AGG GAA GCG TCT ACC TA – 3’

This protocol produces PCR products of 359 bp in wild type mice, 397 bp in *floxed FosB* mice, and both bands in heterozygous mice (Figure 2B). We maintain a colony of these mice by breeding heterozygous animals, and use homozygous floxed and wild type littermate progeny in all experiments.

### Validation of *floxed FosB* mice

In order to determine whether expression of Cre recombinase in *floxed FosB* mice effectively eliminates *FosB* gene expression, we mated *floxed FosB* mice with Rosa26^eGFP-L^10^a^ reporter mice (Jackson Labs, stock #024750). This line produces enhanced green fluorescent protein in the presence of Cre recombinase. We then stereotaxically injected herpes simplex virus expressing Cre (HSV-Cre; MIT Viral Vector Core, Boston, MA, USA) into the hippocampus of adult male wild type and *floxed FosB* mice carrying this reporter as previously described (Eagle et al., 2015). 48 hours later, mice were euthanized by chloral hydrate overdose and transcardially perfused with ice cold 10% formalin to fix brain tissue. Brains were post-fixed in 10% formalin for 24 hours at 4°C, then cryoprotected in 30% sucrose. Coronal sections (35 μm) were prepared on a freezing microtome and then stored in PBS with 0.1% sodium azide. Immunofluorescent labeling was performed using a goat anti-GFP antibody (Abcam, ab5450, 1:1000) and a rabbit anti-FosB antibody (Cell Signaling, 5G4; 1:500), with secondary antibodies from Jackson Immunoreagents.

We observed that HSV-Cre-mediated eGFP had a broad neuronal expression profile throughout the hippocampus in both wild type and *floxed FosB* mice (Figure 2C). In the CA1 region of wild type mice, about 10% of HSV-Cre transduced neurons were found to be FosB-positive, consistent with our previous results (Eagle et al., 2015). However, in *floxed FosB* mice, we found that less than 2% of HSV-Cre transduced cells had a detectable FosB expression, an almost total abrogation of *FosB* gene activity (Figure 3A and B; t=3.603; df=10). These data demonstrate that the *floxed FosB* mouse does not express the *FosB* gene in the presence of Cre recombinase, validating this mouse line for investigation of the role of *FosB* gene products in various cellular and behavioral phenotypes.

**Figure 3:**
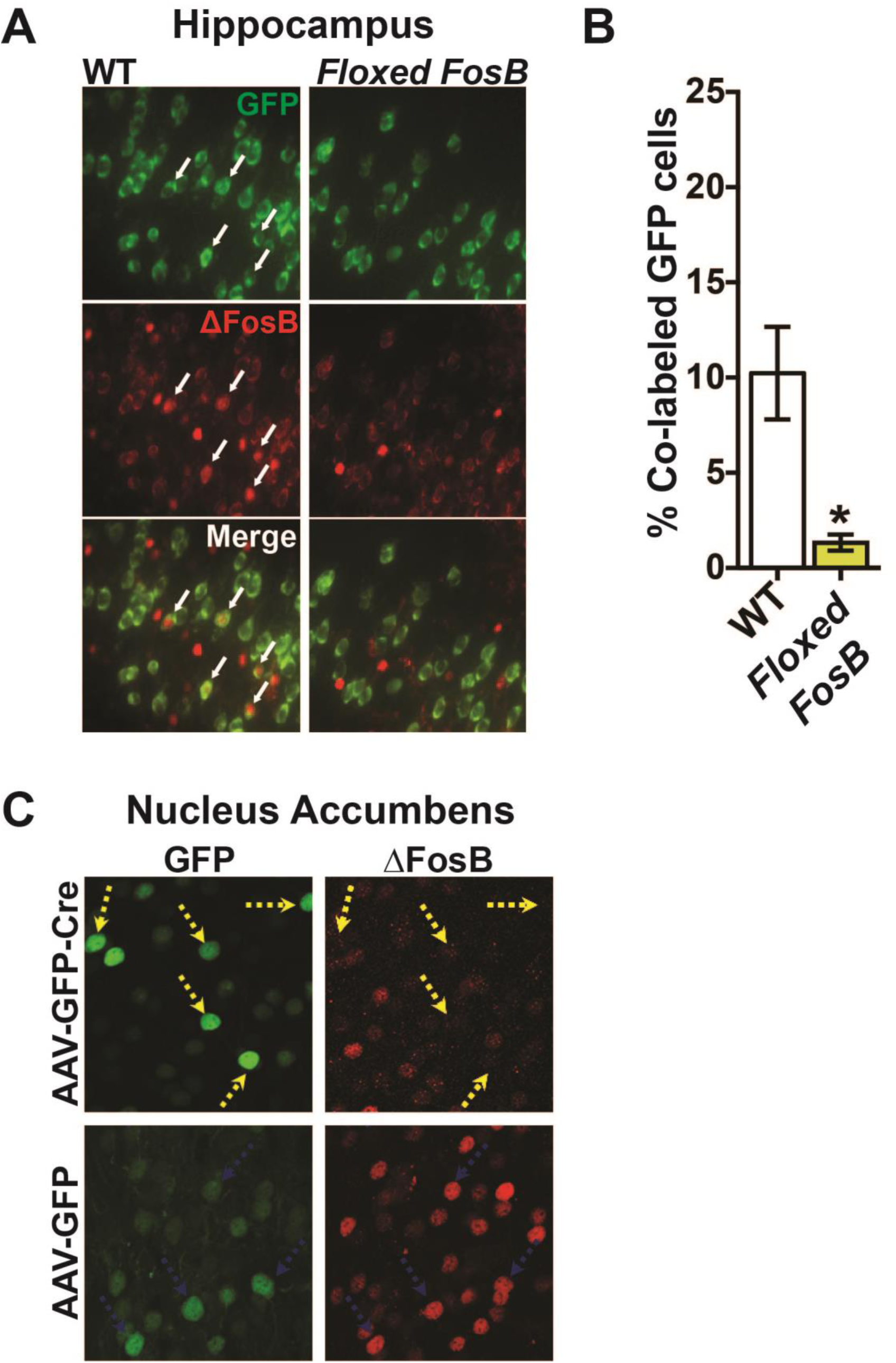
*Floxed FosB* mouse *in vivo* validation. A) Immunofluorescent labeling of GFP expressed in HSV-Cre-transduced neurons (Green) and FosB (red) in hippocampus of wild type (WT) or *floxed FosB* mice. White arrows indicate FosB-positive transduced cells in the WT mouse which are absent in the *floxed FosB* mouse. B) Quantification of data from (A) demonstrating Cre-dependent knockout of FosB protein expression in the *floxed FosB* mouse. Error bars indicate SEM, n=6 mice per group, *:p<0.005 compared to WT. C) Immunofluorescent labeling of GFP expressed in AAV-GFP or AAV-GFP-Cre-transduced neurons (Green) and FosB (red) in nucleus accumbens of *floxed FosB* mice 90 minutes after acute cocaine exposure (i.p., 20 mg/kg). Yellow arrows indicate GFP-Cre-transduced cells lacking FosB, blue arrows indicate GFP-transduced cells which continue to express FosB.

In order to ensure that these results would generalize to other viral methods and brain regions, *Floxed FosB* mice were given intra-NAc injections of AAV-Cre-GFP or AAV-GFP (University of North Carolina Vector Core, Chapel Hill, NC, USA). Three weeks later, when transgene expression is maximal, mice were given a single IP dose of cocaine (20 mg/kg; Sigma Aldrich, St. Louis, MO, USA). Animals were euthanized 90 min later when FosB induction is known to be maximal, and immunofluorescence was performed as above. Induction of FosB was dramatically reduced in animals expressing Cre (Figure 3C).

## Discussion

Here, we describe the production and validation of a *floxed FosB* mouse line, a novel tool for exploration of the role of *FosB* gene products in a variety of contexts. Future experiments may involve mating this line with multiple Cre driver lines to assess the roles of *FosB* gene expression in specific cell types. Of primary interest are driver lines allowing selective *FosB* deletion in D1-type or D2-type medium spiny neurons (Lobo et al., 2013), and lines allowing investigation of the role of the *FosB* gene in hippocampal pyramidal cells (Eagle et al., 2015). As we demonstrate here, the use of viral vectors to express Cre and inhibit *FosB* gene expression in a spatially and temporally restricted manner will also be a viable approach. Moreover, the use of retrograde viruses expressing Cre will allow circuit-specific investigation of the cellular role of *FosB* gene expression.

A critical limitation to investigation of the role of ΔFosB in cell function and behavior has been the limited ability to identify target genes. Early gene expression and microarray analyses revealed many candidates (McClung and Nestler, 2003), but poor antibodies have made unbiased ChIP sequencing experiments difficult. By combining our *floxed FosB* mouse with Cre driver lines and reporter lines allowing expression of GFP-tagged ribosomal subunits, translating ribosome affinity purification (TRAP) (Heiman et al., 2014) may be able to reveal genes whose expression is altered when *FosB* is silenced. Moreover, combining viral GFP-Cre transduction of *floxed FosB* and wild type mice with immunofluorescent detection of expression of candidate target gene products will allow *in vivo*, cell type-specific validation of such targets. Thus, the *floxed FosB* mice presented here represent an exciting new tool for the continued investigation of this critical gene.

